# Evidence of Human adaptation for cluster 2 TMUV strain despite effective anti-viral responses

**DOI:** 10.1101/2025.10.16.682760

**Authors:** Kachaporn Jintana, Takuma Ariizumi, Kanana Rungprasert, Wikanda Tunterak, Julien Pompon, Koshiro Tabata, Aunyaratana Thontiravong, Arnaud Monteil, Rodolphe Hamel

## Abstract

Tembusu virus (TMUV) is an emerging *Orthoflavivirus* with potential zoonotic importance. To date, the biological differences among TMUV strains remain poorly understood. In this study, we compared the replication of a TMUV strain belonging to the cluster 1 and one belonging to the cluster 2 in human lung epithelial cells and further examined the replication characteristics of TMUV cluster 2 strain in both human and mosquito cell lines. The TMUV cluster 2 strain demonstrated a significant higher replication efficiency than the cluster 1 strain in human cells, indicating stronger adaptation to human-derived cells. Infection of human cells with the cluster 2 strain also induced robust innate immune response, including type I interferon and interferon-stimulated gene expression, associated with activation of the unfolded protein response via the PERK, and IRE1 pathways, reflecting an ER stress engagement. These findings suggest that TMUV cluster 2 strain exhibits enhanced replication capacity and more effectively modulates human host responses at the cellular level, underscoring its potential relevance in human health and the zoonotic risk it represents.

## Introduction

Tembusu virus (TMUV) is a mosquito-borne virus that emerged recently and gradually obtained attention following outbreaks in poultry farm in Asia, becoming a major concern in avian health and showing its zoonotic potential (1). TMUV belongs to the genus *Orthoflavivirus* within the family *Flaviviridae* (2) and it was first isolated from *Culex tritaeniorhynchus* in 1995 in Kuala Lumpur, Malaysia, and remained then periodically reported in other country in South-East Asia during the 1970’s (3). Since the first major outbreak of TMUV in 2010 in China, TMUV became an important veterinary pathogen, causing widespread and severe disease outbreaks in egg-laying ducks and spread widely throughout Center Asian and South-East Asian countries (4–8). TMUV was shown to replicate in various organs including the brain, ovary, spleen, lung, reproductive tractus and gastrointestinal tract, leading to internal hemorrhage and triggering encephalitis and neurological disorder in infected bird. Morbidity and mortality rate of infected ducks can reach 80–90%, and 10–30%, respectively (9–11). In addition, TMUV infection is also responsible of a dramatic decrease in egg production named egg drop syndrome, causing significant economic losses in the poultry industry (12). Phylogenetic analyses reveal that TMUV is genetically classified into 4 distinct clusters (cluster TMUV, cluster 1, cluster2 and cluster 3) informed by genetic divergence, with Cluster 1 and Cluster 2 representing the major circulating lineages (1). Notably, airborne transmission has been reported for Cluster 2, highlighting alternative routes of virus spread beyond traditional vectors (13).

TMUV is mainly transmitted through biting of *Culex* genus mosquitoes, however, alternative modes of transmission are also possible including direct contact, fecal-oral route as well as airborne transmission (4, 13, 14). Beside duck infection, TMUV has the ability to infect other birds such as chickens (15, 16), geese (17), sparrows (18) and quails (19) suggesting that TMUV has a wide avian host range. Although TMUV is well known to infect birds, a range of evidence suggests that it may infect mammals and pose a zoonotic risk. Thus, as early as 2010, Tang *et al* reported TMUV seropositive serum from duck farm workers and TMUV RNA positive farmer’s oral swabs (20). Similar seroconversion evidence were reported after a serological survey in human in Thailand. A set of mammalian cells are susceptible *in vitro* to the virus (21). Surprisingly, a recent retrospective study of dolphins that died after showing neurological symptoms, revealed the presence of infectious TMUV in the brain and other tissues (22). Despite the absence of symptoms in human, these previous reports have shown the ability of the virus to infect a range of hosts including human, demonstrating the zoonotic risk of TMUV (15, 18, 20).

Upon infection, flaviviruses stimulate a significant activation of innate immune system which promotes the production of broad range antiviral molecules. Host cells recognize pathogen-associated molecular patterns (PAMPs), including genomic single-stranded RNA and replication intermediates, through pattern recognition receptors (PRRs) including RIG-I–like receptors (RLRs), and Toll-like receptors (TLRs) (23, 24). This triggers the key transcription factors such as IRF3/7 and NF-κB, resulting in the production of type I interferons (IFNs), along with interferon-stimulated genes (ISGs) and proinflammatory cytokine, to establish an antiviral state (25). In avian species, TMUV has been shown to activate innate responses via MDA-5, RIG-I and TLR3 (26). Moreover, TMUV modulate host cellular to promote their replication and operate the unfolded protein response (UPR) pathway leading to viral evasion by impairing antiviral signaling (27, 28).

Given the potential for the airborne transmission and the severe manifestations of disease associated with infection in birds, but even more the risk of zoonotic TMUV infection, we explore in this study the TMUV infection of human lung cell lines by comparing two different clusters of the virus in order to characterize cell tropism, innate immune response and modulation of the UPR pathway along infection. This work aims to elucidate TMUV-host interactions at human cell level and assess the molecular consequences of a potential spillover of TMUV in humans.

## Materials and Methods

### Cells culture

The A549 human cells line (alveolar basal epithelial cells derived from lung carcinoma, ATCC® CCL-185TM) were culture in 75 cm^2^ tissue culture flasks in cell growth medium containing Dulbecco’s modified Eagle’s Minimum Essential Medium (DMEM) (Gibco^TM^, France), 1% penicillin-streptomycin (Gibco^TM^, France) and supplemented with 10% heat-inactivated fetal bovine serum (FBS) (Eurobio Scientific, France). BHK-21 cells line (Fibroblasts cells derived from baby hamster kidneys, ATCC ® CCL-10TM) were culture in 75 cm^2^ tissue culture flasks in cell growth medium containing Roswell Park Memorial Institute Medium (RPMI) 1640 (Gibco^TM^, France), 1% penicillin-streptomycin and supplemented with 10% heat-inactivated FBS. The cells were maintained in a humidified 37^◦^C incubator with 5% CO_2_. *A. albopictus* C6/36 cells were grown at 28°C with 5% CO_2_, growth medium containing DMEM supplemented with 10% heat-inactivated FBS, 1% of non-essential amino-acid (NEAA) (Gibco^TM^, France) and 1% penicillin-streptomycin. When cells reach 80-90% confluence the monolayer cells were sub-passaged by trypsinization following the ATCC instruction.

### Tembusu virus infectious clone construction using Circular Polymerase Extension Reaction (CPER)

Two infectious clones of TMUV (TMUVic) were generated using bacteria-free reverse genetics system by adapting the Circular Polymerase Extension Reaction (CPER) method described by Dong et al (29). The TMUV cluster 1 strain DK/TH/CU-DTMUV2007 (GeneBank accession number MF621927) isolated in 2007 from duck in 2007 and the TMUV cluster 2 strain DK/TH/CU-85 (GeneBank accession number ON931346), isolated in 2016 from duck in Thailand and known for not easily replicating in mosquitoes, were used in this study as template to generate the infectious clones.

The viral RNA from the two strains were extracted using the NucleoSpin RNA Virus extraction kit (MACHEREY-NAGEL, France) and used to generate cDNA by reverse transcription following manufacturer instruction (Promega, France). Viral genome sequences were used as template to design a set of specific primers covering the entire genome within 3 or 4 fragments of the strain TMUV DK/TH/CU-85 and the strain DK/TH/CU-DTMUV2007 respectively (S1 table). The amplification of every fragment was carried out using Q5 High-Fidelity DNA Polymerase (New England Biolabs, France) following manufacturer instruction. A linker DNA fragment was obtained by DNA assembly as follow. The linker comprised the cytomegalovirus (CMV) promoter sequence, the simian virus 40 polyA (SV40 polyA) signal for transcription termination sequence, both obtained using pCDNA3.1 plasmid as template, and the hepatitis delta virus ribozyme sequence (HDVr) and obtained by chemical synthesis (Eurofins Genomics, France). The latest 3 parts were amplified independently by PCR (primer in S1 table) and contained overlapping fragment. All fragments were assembled using NEBuilder HiFi DNA Assembly reaction (New England Biolabs, France) following manufacturer instruction to form the linker fragment. Briefly, equimolar amount at 0.1 pmol of each fragment were pooled and mixed with NEBuilder HiFi DNA Assembly Master Mix and Nuclease free water in total volume of 20 µL and incubate at 50°C for 15 minutes. Purified TMUV DNA amplicon fragments and the linker DNA fragment construct contain overlapping sequences for annealing assembly during the CPER. For each strain, TMUV DNA amplicon fragments and the linker DNA fragment were mixed in equimolar amounts in PCR reaction containing 10 µL of Q5 Buffer (New England Biolabs, France), 1 µL of 10mM dNTPs (Thermo Fisher Scientific, France) and 0.5µ L of Q5 high fidelity polymerase (New England Biolabs, France) for a final volume of 50µL. The following cycling conditions were used: initial denaturation at 98°C for 2 min, followed by 30 cycles of denaturation at 98°C for 30 s, and annealing/extension step at 68°C for 8 min and a final extension at 68°C for 10 min. Product from the latter CPER assembly were then transfected into BHK-21 cells line in 12-well plates using lipofectamine 2000 (Thermo Fisher Scientific, France) as a transfection reagent. Supernatants containing TMUV infectious viral clone (TMUVic) progeny were harvested at 6 days post-transfection and cells were fixed using 3.7% paraformaldehyde in PBS for IF assay (described below). Viral RNA was extracted from the supernatant using the NucleoSpin RNA Virus extraction kit (MACHEREY-NAGEL, France) according to the manufacturer’s instructions. Viral RNA copy number were determined by RT-qPCR following the method described below. TMUVic where then amplified from the supernatant collected at day 6 in 75 cm^2^ tissue culture flasks to generate a large batch of virus. Titers of the virus production were determined by focus forming assay (FFU) described below.

### Virus titration by focus forming assay (FFU)

Confluent BHK-21 cells monolayer was prepared in 24-well tissue culture plates for 24 h. The supernatant containing TMUVic was serial 10-fold diluted in mixed RPMI 1640 medium supplemented with 1% Penicillin-Streptomycin. Each concentration of TMUVic mixed was transferred to monolayer BHK-21 cells. The inoculum cells were incubated at 37^◦^C with 5% CO_2_ for 1 h. with gently rock the plate on a rocker for viral adsorption. At the end of incubation time, inoculum was gently discarded and infected in each well were overlayed with 2% Carboxymethylcellulose (CMC) (Sigma-Aldrich, France) in FBS-RPMI 1640 supplemented with 1% of Penicillin-Streptomycin, then incubated at 37^◦^C with 5% CO_2_ for 4 days. Each well was fixing with a 3.7% paraformaldehyde solution and fixing cells were incubated with Triton X-100. Foci were detected using mouse anti-flavivirus group antigen clone D1-4G2-4-15 (4G2) antibody (Sigma-Aldrich, France) with goat anti-mouse IgG secondary antibody Alexa Fluor^TM^ 488 (Sigma-Aldrich, France). The number of foci were count under the fluorescent microscope (EVOS M5000 imaging system, Invitrogen, France) and virus titer was calculated from the average of triplicate wells and presented as focus forming unit per milliliter (FFU/mL).

### Virus immunofluorescence staining assay (IFA)

To visualize the infection of the different cell lines used in this study, infected cells were fixed with 3.7% paraformaldehyde in PBS for 20 min at room temperature and permeabilized using 0.5% Triton X-100 solution in PBS for 10 min at room temperature. Cells were blocked with a 10% FBS and 0.3% Triton X-100 solution treatment for 30 min at 37°C then incubated for 1 h at 37°C with the monoclonal antibody (MAb) 4G2 mouse anti-flavivirus group antigen clone D1-4G2-4-15 (Sigma-Aldrich, France). Cells were washed 3 times with 1X PBS and incubated for 1 h at 37°C with goat anti-mouse IgG secondary antibody conjugated to Alexa Fluor^TM^ 488 (Invitrogen, France). ProLong^TM^ Gold antifade reagent with DAPI (Invitrogen, France) was used as mounting medium and nucleus counterstain dye. Fluorescent images were generated under the EVOS M5000 imaging microscope system (Invitrogen, France).

### Viral RNA extraction and quantitation by one step Reverse Transcription real-time polymerase chain reaction (RT-qPCR)

Viral RNA was extracted using the NucleoSpin RNA Virus extraction kit (MACHEREY-NAGEL, France) according to the manufacturer’s instructions. The viral RNA of TMUV was quantitated by RT-qPCR using the GoTaq^®^ 1-Step RT-qPCR System (Promega, France). The primer sequences targeting TMUV are listed in S2 Table (30), RNA copies were used to construct a standard curve. and reaction was performed using the LightCycler^®^ 96 (Roche, Switzerland). Viral RNA was quantified by comparing sample’s threshold cycle (CT) value with a standard curve made with a serial dilution of a known concentration of TMUV RNA. The LightCycler^®^ software was used to analyze the results and determine viral RNA concentration in copies/mL.

### RNA preparation

Total RNA from cells culture which were infected with TMUVic was extracted using Tri-Reagent (Euromedex, France) according to the manufacturer’s protocol. Briefly, cells were homogenized in Tri Reagent and incubated for 5 min at RT. Chloroform was then added, and the mixture was vigorously shaken and centrifuged at 12,000 g for 15 min at 4°C to separate the phases. The aqueous phase containing RNA was carefully transferred to a new microcentrifuge tube, and total RNA was precipitated by the addition of isopropanol followed by centrifugation at 12,000g for 8 min at 4°C. The supernatant was removed, and RNA pellet was washed twice in 75% ethanol and subsequently centrifuge at 7,500g for 5 mins at 4°C. Finally, RNA pellet was air-dried, and resuspended in nuclease-free water. RNA concentration and purity were assessed using a NanoDrop^TM^ One spectrophotometer Microvolume UV-Vis Spectrophotometer (Thermo scientific, France).

### Reverse transcription and real time PCR for gene expression investigation

A quantity of 1 µg of total RNA from cell samples was reverse transcribed into complementary DNA (cDNA) using the M-MLV Reverse Transcriptase kit (Promega, France). The reaction mixture contained random primer (Promega, France) RNaseOut^TM^ (Invitrogen, France) and a set of dNTP (Euromedex, France) in final reaction volume of 20 µL. Reverse transcription was performed under the following conditions 37°C for 60 min, 85°C for 10 min. The cDNA samples were stored at −20°C until use for the real time PCR.

Real-time PCR was carried out for a set of selected genes using specific primers listed in S3 Table. Real-time PCR was performed using the 5x Hot Pol EvaGreen^TM^ qPCRreaction master mix (Euromedex, France) under the following conditions: a first activation step at 95°C for 10 min, followed by 45 cycles of 95°C for 15 s, 60°C for 15 s and 72°C for 20 s. A melting curve was generated at the end of the run. The mRNA relative expression (fold change) was calculated using the 2^−ΔΔCT^ method with comparison of the threshold cycles (CTs) for each target gene of interest with the CT of the GAPDH gene used as the endogenous control.

### PCR analysis for XBP-1

The cDNA sample were amplified by PCR to analyze XBP1 using specific primer (S3 Table). Amplification conditions included an initial denaturation at 95°C for 5 min, followed by 45 cycles of 95°C for 30 s, 65°C for 30 s, and 72°C for 30 s, and a final extension at 72°C for 5 min. PCR products were resolved by electrophoresis in a 2% agarose gel stained with GelRed^®^ Nucleic Acid Gel Stain (Biotium, France).

### RNA interference and gene overexpression

Cells were prepared in 48 well plates and transfected using lipofectamine 2000 (Invitrogen, France) with 5 pmol per well of specific siRNAs targeting MDA transcript (flexitube ref#SI04765852, Qiagen, France) or TLR3 transcript (flexitube ref#SI02630782, Qiagen, France) or with a non-silencing siRNA (allstars multiplex siRNAs ref#SI91027281, Qiagen, France).

Cells were prepared in 48 well plates and transfected using lipofectamine 2000 (Invitrogen, France) with 5 µg of each plasmid encoding MDA5, RIG-I, TLR3, or TLR7 (Addgene, USA) according to manufacturer’s instruction.

After the 24 hours, cells transfected with siRNAs or plasmids were infected with TMUVics at MOI 0.1. The supernatant and cells were collected every day until 4 days of post infection in treated for gene expression and virus replication quantitation as mentioned above.

### Statistical analysis

Data are presented as the mean ± SEM from three independent experiments. The significance of the differences groups was evaluated by using one-way and two-way ANOVA analysis followed by a multiple comparison test as appropriate. Graphs were plotted and analyzed using the GraphPad Prism software version 10. A p-value less than 0.05 was considered statistically significant.

## Results

### Generation of TMUV infection clone by CEPR approach

To investigate the possibility of TMUV adaptation to vertebrate hosts, we generated two infectious viral clones corresponding to two TMUV from the cluster 1 and the cluster 2, using reverse genetics approach. As described by Aubry et al. (31) reverse genetics facilitates the study of numerous aspects of host-flavivirus interaction mechanisms. Such approaches enable the production of genetically homogeneous viral clones, thereby overcoming the variability that can be found in strains obtained from field samples. To generate our clones, we drew inspiration from the circular polymerase extension reaction (CEPR) technique previously published by Dong *et al* (29). For the TMUV cluster 1 (TMUVic-C1), we generated by PCR four overlapping sequences covering the entire genome, then generated circular DNA by CEPR, which we transfected into BHK21 cells (Fig 1A). For TMUV cluster 2.1 (TMUVic-C2), we generated by CEPR the circular DNA with three overlapping sequences, which we also transfected into BHK21 cells.

**Fig 1.**
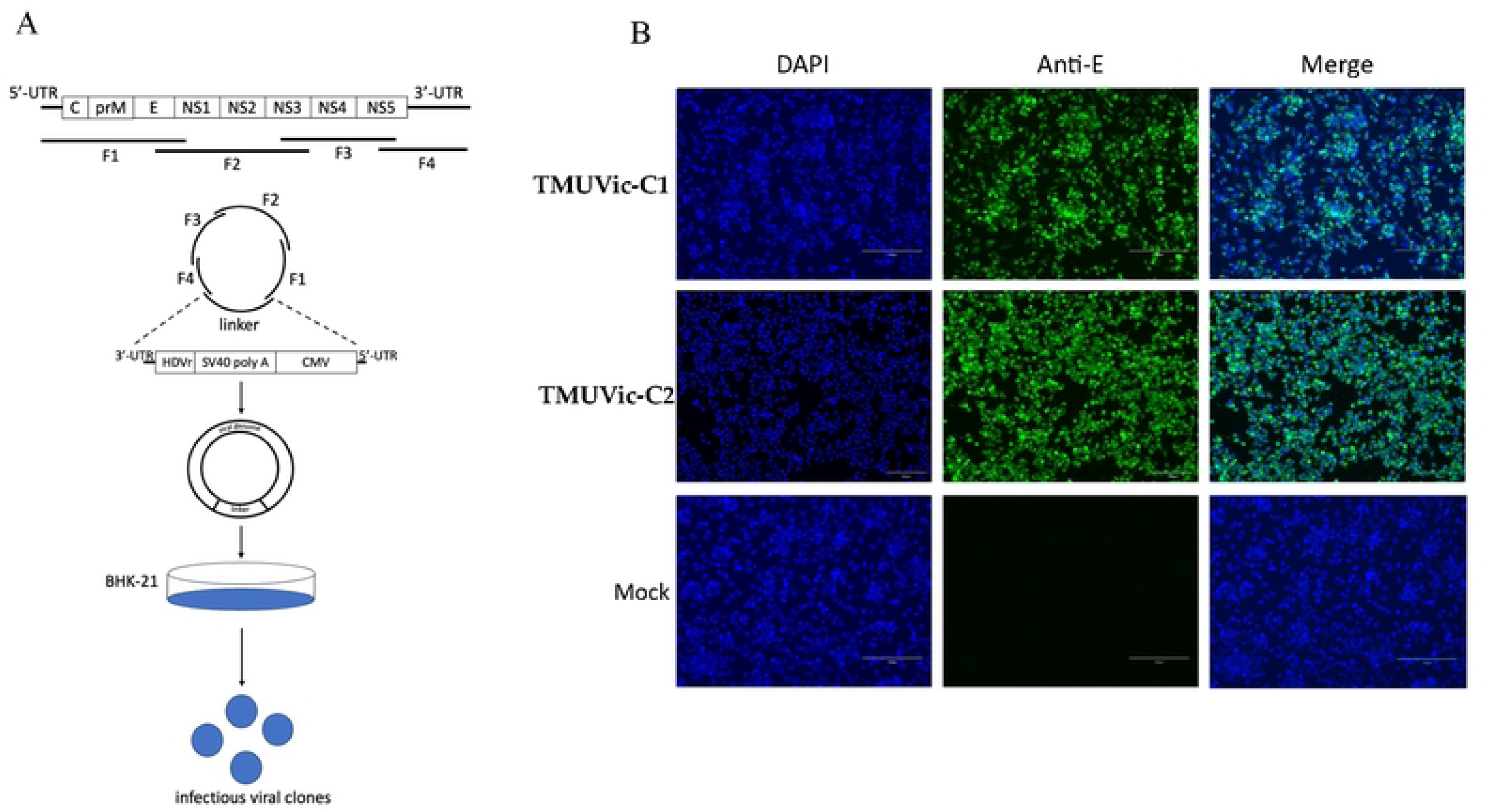
Development of TMUV infectious clone (TMUVic) using CPER approach. (A) Scheme for assembly of TMUV genome cluster 1 (4 fragments F1 to F4) and cluster 2 (3 fragments F1 to F3, not shown on the diagram) using CPER method and BHK21 cell line to recover infectious virions. (B) Confirmation of infectivity of TMUVic-Cluster-1 (TMUVic-C1) and -Cluster-2 (TMUVic-C2) using IFA at 6 days post infection (dpi). E protein expressed by TMUVic was identified using IFA analysis with a monoclonal 4G2 antibody against Flavivirus E protein (green). Cell nuclei were stained with DAPI (blue).

Six days after transfection, we detected viral envelope proteins in the majority of cells from both batches transfected with the TMUVic-C1 and TMUVic-C2 construct, unlike the mock-transfected cells, which showed no staining with the anti-viral envelope protein antibody (Fig 1B). This observation showed the effective replication of each viral clones in BHK21 and confirmed that we obtained two infectious and replicative viral clones belonging to Cluster 1 and Cluster 2.

### Cluster-specific TMUV infectivity in Human lung cells

Given previous studies reporting the ability of TMUV of the cluster 2 to infect avian hosts via the airborne route, we wonder if TMUV cluster 2 strain can infect human cell from respiratory tract compared to a TMUV strain known to be an arthropod born strain. To test this hypothesis and evaluate whether human lung epithelial can serve as entrance for TMUV in humans, we first tested the permissiveness of the A549 – human lung epithelial cells to be infected by TMUVic-C1 (derived from the cluster 1 strain) and TMUVic-C2 (derived from strain and cluster 2). Cells were infected with each virus at a MOI of 0.1 and the viral replication was monitoring over 3 days by quantification of the viral RNA genome. We reported a robust viral replication of the TMUVic-C2, reaching 10^6^ copies/µL after 72 hpi, while a non-significative increase in viral copy number was denoted with TMUVic-C1 (Fig 2A). Same striking differences were observed between the two clusters regarding viral titer with a concentration 10^4^ FFU/mL of TMUVic-C2 at 3 days post-infection (dpi), while TMUVic-C1 strain failed to establish productive infection in A549 cells (Fig 2B). The ability of A549 to support TMUVic-C2 replication over a long period of time was then investigated. Infection of A549 resulted in a gradual increase in viral copy number in cell supernatant as well as in the intracellular compartment, which peaked at 4 days after infection, followed by a very slight decline to a plateau at 7 days (Fig 2C and 2D). This replication profile is confirmed by measuring the production of viral progeny in cell supernatant by focus forming assay (Fig 2E). These observations demonstrate a cluster-specific difference in human cell tropism, with Cluster 2 virus having the ability to replicate efficiently in human lung epithelial cells, unlike the Cluster 1 virus.

**Fig 2.**
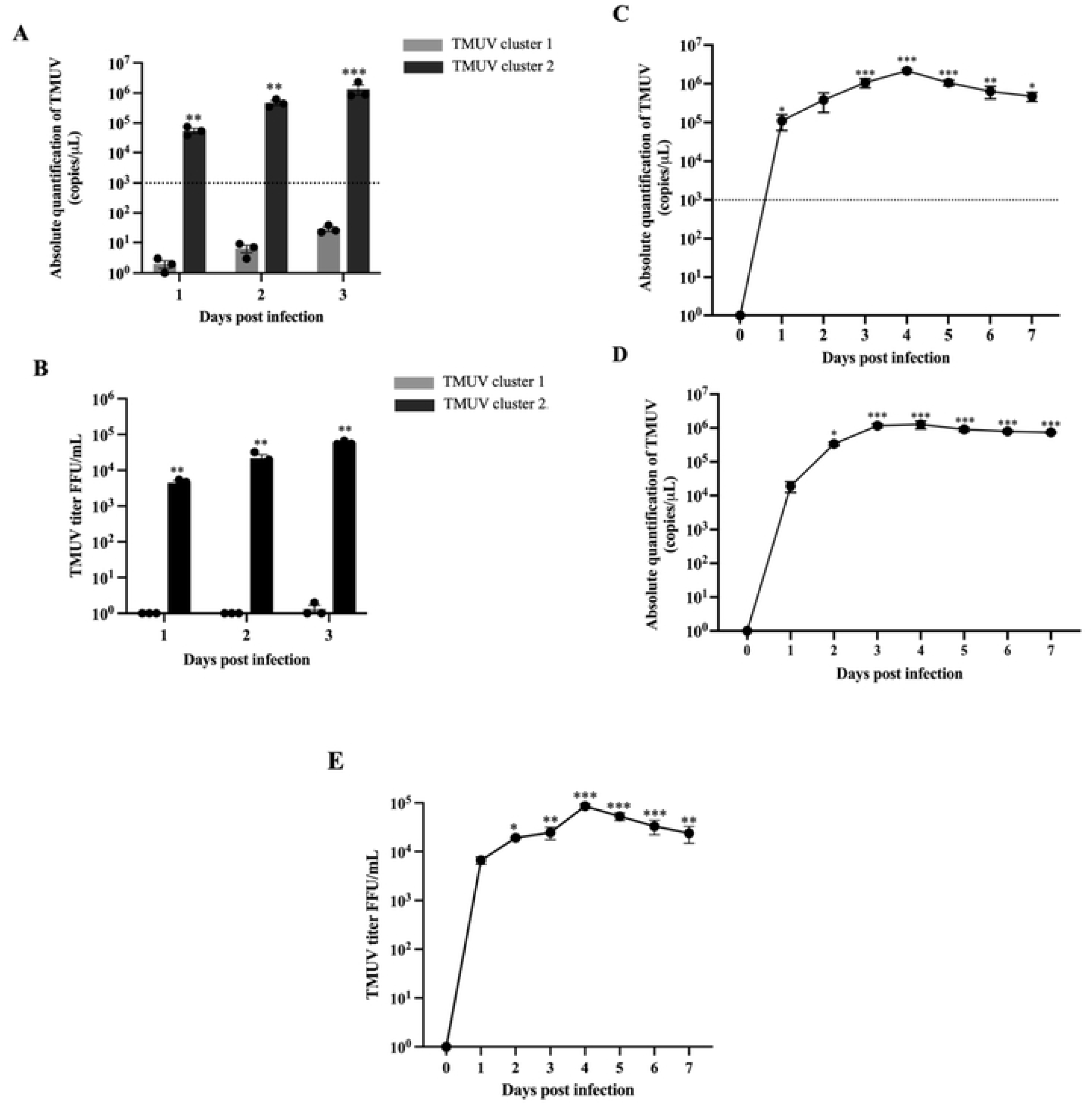
Differential replication capacity of TMUV genetic clusters in human lung epithelial cells. Cells A549 were infected at a MOI of 0.1 with both TMUVic-C1 and TMUVic-C2 strains and viral replication was monitored over 3 days post-infection (dpi). (A) Expression of viral RNA was quantified by RT-qPCR in RNA copies/µL B) Viral replication was measured using focus-forming assay (FFA) and expressed as FFU/mL. (C) Kinetics of TMUVic-C2 infection in human lung epithelial A549 cells. Quantification of TMUVic-C2 copy number in supernatant of infected cells. (D) Quantification of TMUVic-C2 copy number in internal compartment of infected cells. (E) Viral replication of TMUVic-C2 was measured using focus-forming assay (FFA). Data represent mean ± SEM of three independent experiments. *\*\*p < 0.01, ***p < 0.001* compared to A549 cells infected TMUVic-C1.

### TMUV cluster 2 can maintain a more active replication in human lung cells as compared to mosquito cells

Knowing that cluster 2 is preferentially aerially transmitted and infects human cells, we wondered whether this strain was well-adapted to replicate efficiently in mosquitoes. Viral replication potential of TMUVic-C2 was compared between the A549 human lung epithelial cells and the C6/36 mosquito cells derived from *Aedes albopictus* (32). Both cell types were infected with TMUVic-C2 at a multiplicity of infection of 0.1 and monitored for a 4-day period. Quantification of viral RNA by RT-qPCR and viral replicative particles by FFA revealed that A549 cells exhibited significantly higher levels of TMUVic-C2 as compared to C6/36 cells (Fig 3A and 3B). In A549 cells, viral RNA levels showed an exponential increase up to 4 dpi, reaching more than 10^6^ viral RNA copies/μl (Fig 3A) and a titer of over 10^5^ FFU/mL (Fig 3B). In contrast, C6/36 cells displayed a maximal viral RNA copies levels less than 10^3^ copies/μl which is the minimum threshold value of acceptable quantification accuracy under our experimental conditions (Fig 3A) and a viral particles titers less than 10 FFU/mL (Fig 3B). Active viral replication was confirmed by immunostaining with antibody targeting the viral envelope protein. No staining was observed at 4 dpi in the uninfected control sample while, as soon as 1 dpi, the viral envelope protein was gradually detected in cells up to 4 dpi demonstrating a time-dependent increasing of the virus replication in fluorescence (Fig 3C). Finally, these results show that A549 cells are more permissive to TMUVic-C2 infection as compared to insect cells and may serve as a suitable *in vitro* model to investigate TMUV pathogenesis in human respiratory epithelium.

**Fig 3.**
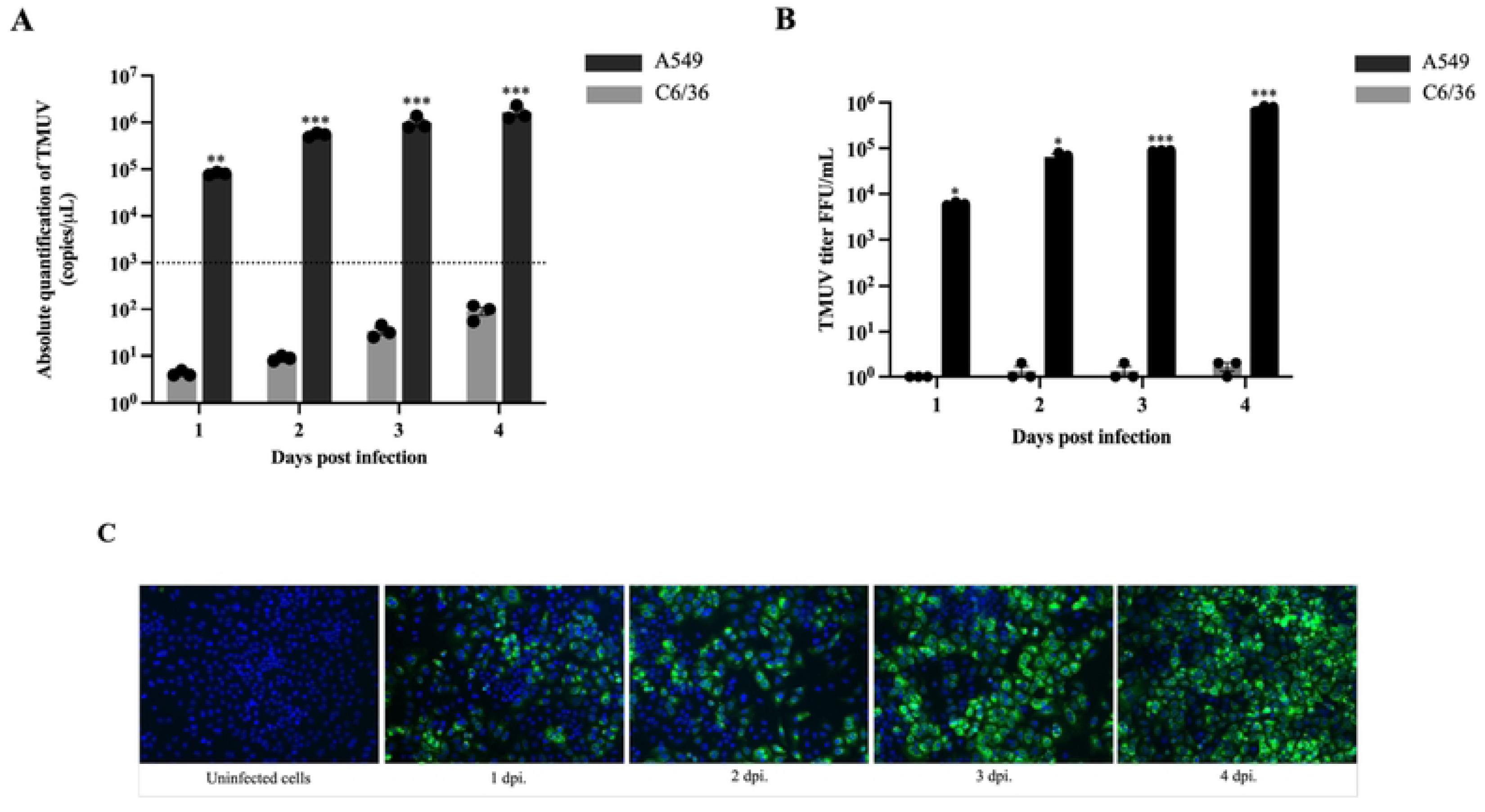
Differential permissiveness of human and mosquito cells to TMUV. A549 human lung epithelial cells and C6/36 mosquito-derived cells were infected with TMUVic-C2 at a MOI of 0.1. Viral replication was monitored over a period of 4 days. (A) Quantification of viral RNA levels at indicated time points and expressed in RNA copies/μL. (B) Infection intensity was determined by plaque assay of culture supernatants of cells exposed to TMUVic-C2. (C) Human lung epithelial cells infected with TMUV at MOI of 0.1 were analyzed at different day post infection for the presence of the viral envelope protein by immunofluorescence with the 4G2 MAb and an Alexa Fluor^TM^ 488-conjugated anti-mouse IgG. Data represent mean ± SEM of three independent experiments. *\*p < 0.05, **p < 0.01, ***p < 0.001*.

### TMUV cluster 2 induces an innate immune response in human lung epithelial cells

To determine whether TMUV cluster 2 strain induces an innate antiviral immune response in permissive human lung epithelial cells, the antiviral transcriptomic profile was determined along the replication cycle of the virus. The key innate immune response genes involved, including cytokines and interferon-stimulated genes (ISGs), were analyzed in A549 cells at various time points post-infection using quantitative RT-PCR. Cells were infected with TMUVic-C2 at a MOI of 0.1, the cells were harvested at 12h dpi then daily until 7 dpi. Mock uninfected cells were used as control. Pattern recognition receptors (PRRs) involved in viral RNA sensing were rapidly activated in response to TMUV infection. Thus, both Toll-like receptors (TLRs) and Retinoic acid-inducible gene I (RIG-I)-like receptors (RLRs) were modulated. A significant up-regulation of RIG-I expression by 12 hpi was observed, peaking at 24 hpi with a 600-fold induction (p < 0.001) and subsequently a marked downregulation of 2.4 times at 72 hpi (Fig 4A). MDA5 showed a dissimilar trend, gradual increase of expression was observed over time reaching a maximum of 109-fold at 96 hpi (Fig 4B). Despite down-regulation in the first hours of the infection, TLR3 was highly upregulated reaching a plateau at 5 dpi (Fig 4C), while TLR7 was moderately increased after 5dpi (Fig 4D). These results suggest that TMUV RNA is sensed by both cytoplasmic and endosomal PRRs.

**Fig 4.**
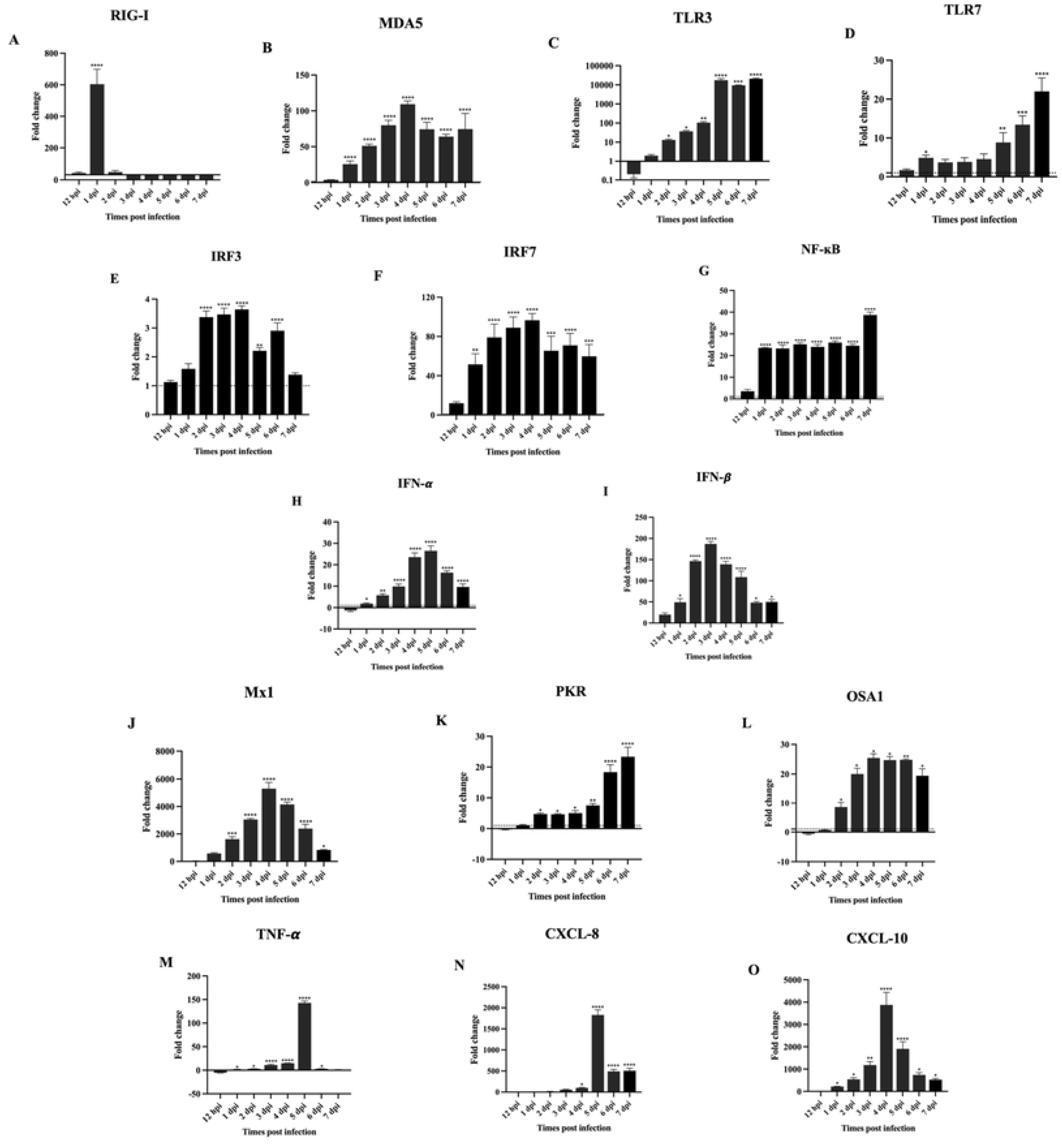
Innate immune responses profile in human lung epithelial A549 cells trigger by TMUV infection. A549 cells were infected with TMUVic at a MOI of 0.1 and harvested at multiple time points from 12 hpi. to 7 dpi. Uninfected cells served as negative controls. The expression of PRRs, adaptor molecules, transcription factors, type I IFN, ISGs, and pro-inflammatory cytokines/chemokines was quantified by RT-qPCR serve as fold change and GAPDH served as housekeeping gene. (A-D) cytoplasmic and endosomal PRRs sensing profile (A) RIG-I, (B) MDA5, (C) TLR3 and (D) TLR7. (E-G) PRR signaling led to activation of transcription factors (E) IRF3, (F) IRF7 and (G) NF-κB. (H-I) Type I IFN responses (H) IFN-α and (I) IFN-β. (J-L) ISGs profile (J) Mx1, (K) PKR and (L) OSA1. (M-O) inflammatory response genes (M) TNF-α, (N) CXCL-8 and (O) CXCL-10. Data represent mean ± SEM of three independent experiments. Statistical significance was determined using one-way ANOVA with post hoc tests (*\*p < 0.05, **p < 0.01, ***p < 0.001* compared to uninfected cell control).

Activation of PRRs by detection of viral PAMPs was accompanied by a modulation of the expression of downstream adaptor molecules and transcription factors particularly involved in signaling pathway. Therefore, IRF3 mRNA level was upregulated lightly after 2 dpi (Fig 4E), while IRF7 and NF-κB expression increased strongly since 1 dpi maintaining over-expression up to 7 dpi (Fig 4F and 4G). This delayed induction implies the activation of signaling pathways essential for interferon and inflammatory gene transcription. To evaluate the latter cascade, we measured the modulation of expression of a set of genes. A robust induction of type I IFN was observed with a significant up-regulation after 1 dpi peaking at 5 dpi (Fig 4H). In a more pronounced way, IFN-β expression levels increasing by 1 dpi with a peak of expression at 3 dpi (Fig 4I). Slightly out of sync with IFN modulation, a marked elevation of interferon-stimulated genes (ISGs) was observed after infection. Expression of Mx1 gene show a strong up-regulation reaching a fold change of 5.298 at 4 dpi before to decreasing to 23.3 at the end of the observation period (Fig 4J). PRK expression increased significantly but mostly at the end of the time course (Fig 4K), while OAS1 showed significant induction with a plateau after 3 dpi (Fig 4L). All together this profile of modulation probably denotes the activation of the JAK-STAT signaling pathway downstream of type I IFN signaling during TMUV infection. In parallel, TMUV infection triggered a pronounced pro-inflammatory response in A549 cells. TNF-α expression was significantly upregulated at 1 dpi (∼3.6-fold) and peaked at 5 dpi with 75.9-fold increase (p < 0.001) (Fig 4M). At the same time, a strong induction of the key chemokines, CXCL-8 and CXCL-10, were observed after 4 and 5 dpi, respectively (Fig 4N and 4O). Taken together, these findings collectively reveal that TMUV activates a comprehensive innate immune response in human lung epithelial cells, involving PRR-mediated sensing, interferon production, ISGs expression, and inflammatory cytokine induction. This complex host response likely plays a critical role in controlling early viral replication but may also contribute to immunopathology during infection.

### PRRs promote innate immune responses against TMUV infection in human lung epithelial cells

To investigate the role of TLRs and RLRs family in antiviral responses against TMUV, a set of PRRs were overexpressed in A549 cells. The validation of the overexpression level of MDA5, RIG-I, TLR3 and TLR7 was done after transfection of A549 cells with specific expression plasmids (S1 Fig). Transfected A549 cells were infected with TMUVic-C2 at a MOI of 0.1 and viral replication was determined by quantitative PCR at 24, 48, and 72 hpi. Compared with the control group, cells overexpressing MDA5, RIG-I, TLR3 or TLR7 exhibited a marked reduction in the viral RNA copy numbers with a more pronounced effect at 48 hpi and 72 hpi (Fig 5A). These findings indicate that both RLRs and TLRs families participate in a strong antiviral activity against TMUVic-C2. Then, we examined the expression of key innate immune genes after infection with TMUVic-C2 of cells transfected with plasmids encoding MDA5, RIG-I, TLR3, or TLR7 (Fig 5B, 5C, 5D, and 5E respectively). The expression of the set of genes was measured at 24, 48, and 72 hpi by qRT-PCR. Infected cells overexpressing of MDA5 present a strong enhancement of type I IFN responses, with IFN-β showing markedly increased induction, peaking at approximately 50-fold above baseline at 48 hpi. IFN-α, IRF3, IRF7, and NF-κB were also significantly upregulated, indicating robust activation of downstream antiviral signaling pathways. Similarly, in RIG-I overexpressing infected cells, type I IFN production is triggered, especially for IFN-β. In addition, IRF7 expression increased significantly, suggesting that RIG-I primarily activates the IRF7-dependent interferon arm. Antiviral signaling is also strongly activated in cells overexpressing TLR3. Both IFN-α and IFN-β were significantly induced, with peak expression observed at 48 hpi for both types, while upregulation in IRF3, IRF7, and NF-κB were more modest. Infection in cells that overexpressed TLR7 presented a significant induction of the expression of type I IFN (IFN-α and IFN-β) as well as the transcription factors IRF3, IRF7, and NF-κB, although to a lesser extent than in cells overexpressing MDA5, RIG-I, or TLR3.

**Fig 5.**
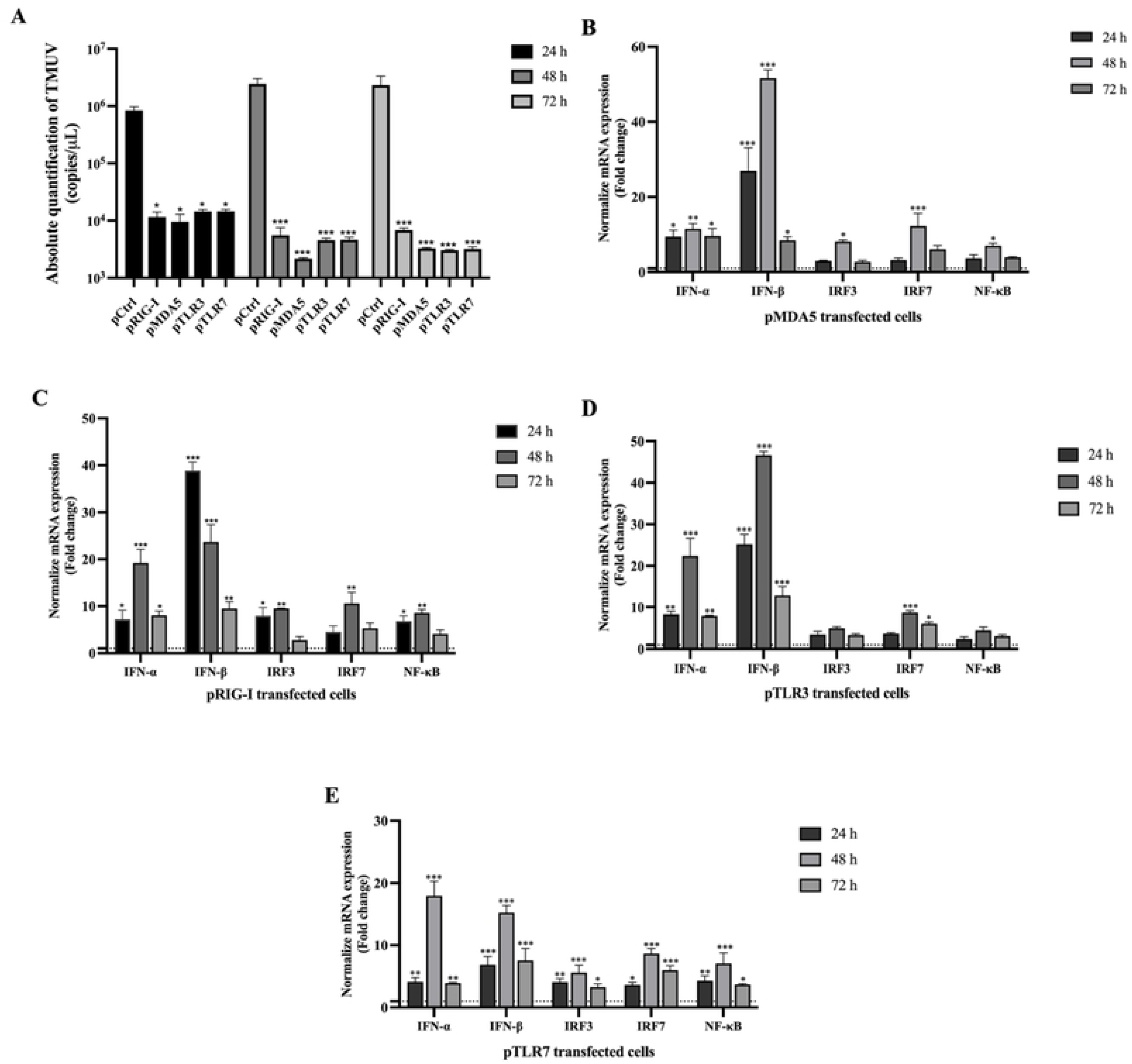
Overexpression of RLRs and TLRs enhances antiviral responses against TMUV in A549 cells. A549 cells were transfected with plasmids encoding MDA5, RIG-I, TLR3, or TLR7 and subsequently infected with cells overexpressing MDA5 (B), RIG-I (C), TLR3 (D), or TLR7 (E) and infected with TMUVic at a MOI of 0.1. (A) Viral replication was assessed by qRT-PCR at 24, 48, and 72 hpi. (B–E) Expression of innate immune genes (IFN-α, IFN-β, IRF3, IRF7, NF-κB) was analyzed in cells overexpressing MDA5 (B), RIG-I (C), TLR3 (D), or TLR7 (E) and infected with TMUVic-C2. Data represent mean ± SEM from three independent experiments. Statistical significance was determined using two-way ANOVA with Tukey’s multiple comparisons tests *(*p < 0.05, **p < 0.01, ***p < 0.001* compared to control).

From previous results, we examined whether reducing receptor expression would conversely promote infection. We performed expression knockdown of MDA5 and TLR3, the two receptors that had shown the strongest antiviral activity in the overexpression assays, using siRNA. Effective knockdown was first validated with either siRNA targeting MDA5 transcript or siRNA targeting TLR3 transcript in A549 cells (S2 Fig). Then, siRNA treated cells were infected with TMUVic-C2 at a MOI of 0.1 and both viral replication and innate immune gene expression were analyzed at 24, 48, and 72 hpi. Firstly, MDA5 gene silencing significantly enhanced viral replication at 24, 48, and 72 hpi, resulting in approximately 1–2 log higher viral RNA levels compared to control cells (Fig 6A). Similarly, knockdown of TLR3 expression also led to increased TMUV replication over time (Fig 6B). Knockdown of MDA5 significantly suppressed the expression of IFN-α, IFN-β, IRF3, IRF7 compared to the control siRNA-transfected cells, while NF-κB showed significant downregulation at later stages (Fig 6C). Similarly, TLR3 knockdown markedly impaired the expression of IFN-α and IFN-β at all time points (Fig 6D). Transcription factors IRF3, IRF7, and NF-κB were also significantly suppressed, underscoring the critical role of TLR3 in initiating and sustaining interferon-mediated antiviral responses. The effect of TLR3 silencing mirrored that of MDA5, confirming that both receptors play essential roles in restricting viral replication and loss of either receptor leads to broad suppression of antiviral gene expression.

**Fig 6.**
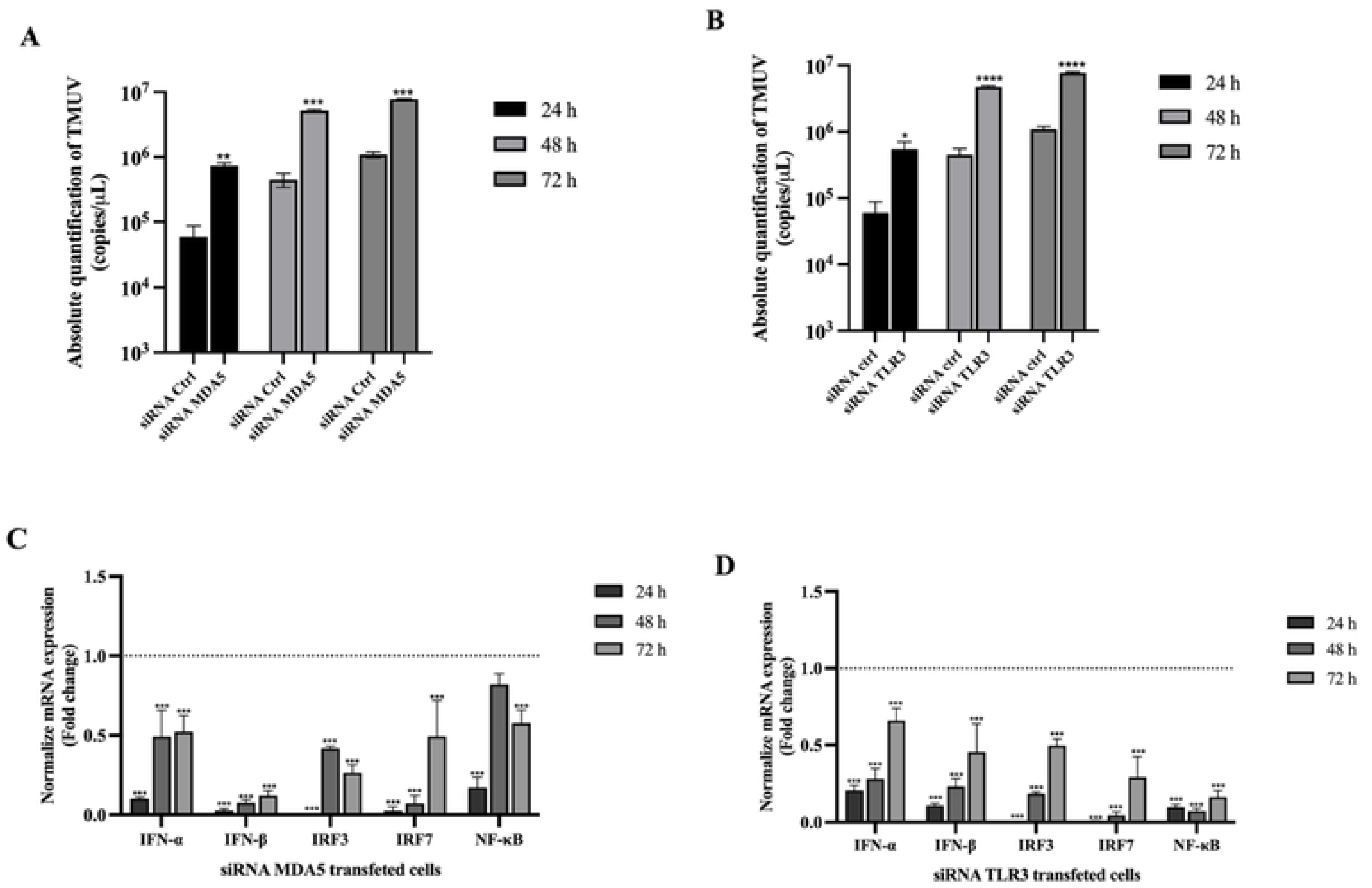
Knockdown of MDA5 and TLR3 expression enhances TMUV replication and impairs antiviral immune responses in A549 cells. A549 cells were transfected with siRNA targeting MDA5 or TLR3 transcript and subsequently infected with TMUVic2 MOI of 0.1. (A, B) Viral RNA levels were quantified by qRT-PCR at the same time points after silencing of MDA5 (A) or TLR3(B). (C, D) Expression of IFN-α, IFN-β, IRF3, IRF7, and NF-κB gene was assessed at 24, 48, and 72 hpi by qRT-PCR after silencing of MDA5 (C) or TLR3(D). Data represent mean ± SEM of three independent experiments. Data represent mean ± SEM from three independent experiments. Statistical significance was determined using two-way ANOVA with Tukey’s multiple comparisons tests *(*p < 0.05, **p < 0.01, ***p < 0.001* compared to siRNA control).

Taken together, these data demonstrate that both RIGI, MDA5, TLR7 and TLR3 are critical PRRs to recognize TMUV infection. These receptors promote a robust innate immune response in human lung cells and underscore their importance against TMUV infection through the induction of type I IFN and key transcription factors.

### TMUV infection induces the activation of the unfolded protein response (UPR) pathway

Besides innate immune response, we also investigated unfolded protein response (UPR) which is activated by endoplasmic reticulum (ER) stress during viral infection and critical for cellular defense against viral infection (33). The UPR is mediated via three domain signaling mechanisms PERK (PKR-like ER kinase), IRE1 (inositol-requiring enzyme 1), and ATF6 (activating transcription factor 6) leading to apoptosis and autophagy (34). To determine whether TMUV infection induces ER stress and activates the UPR in human A549 lung epithelial cells, we analyzed the expression of key UPR-related genes after infection with TMUVic-C2 at a MOI of 0.1. Real time PCR revealed a significant modulation in the expression of at set of UPR-related genes. Among these, CHOP (C/EBP homologous protein), a pro-apoptotic marker of the PERK and ATF6 pathways, was notably up regulated along the time up to a peak at 7 dpi (Fig 7A). We also observed a significant induction of GADD34, a feedback regulator of the PERK-eIF2α pathway, concomitantly to CHOP activation (Fig 7B), and the up-regulation of ATF3, a transcription factor associated with stress response and apoptosis (Fig 7C). These observations indicate the activation of the PERK pathway, which can lead to the activation of ER stress-mediated apoptosis.

**Fig 7.**
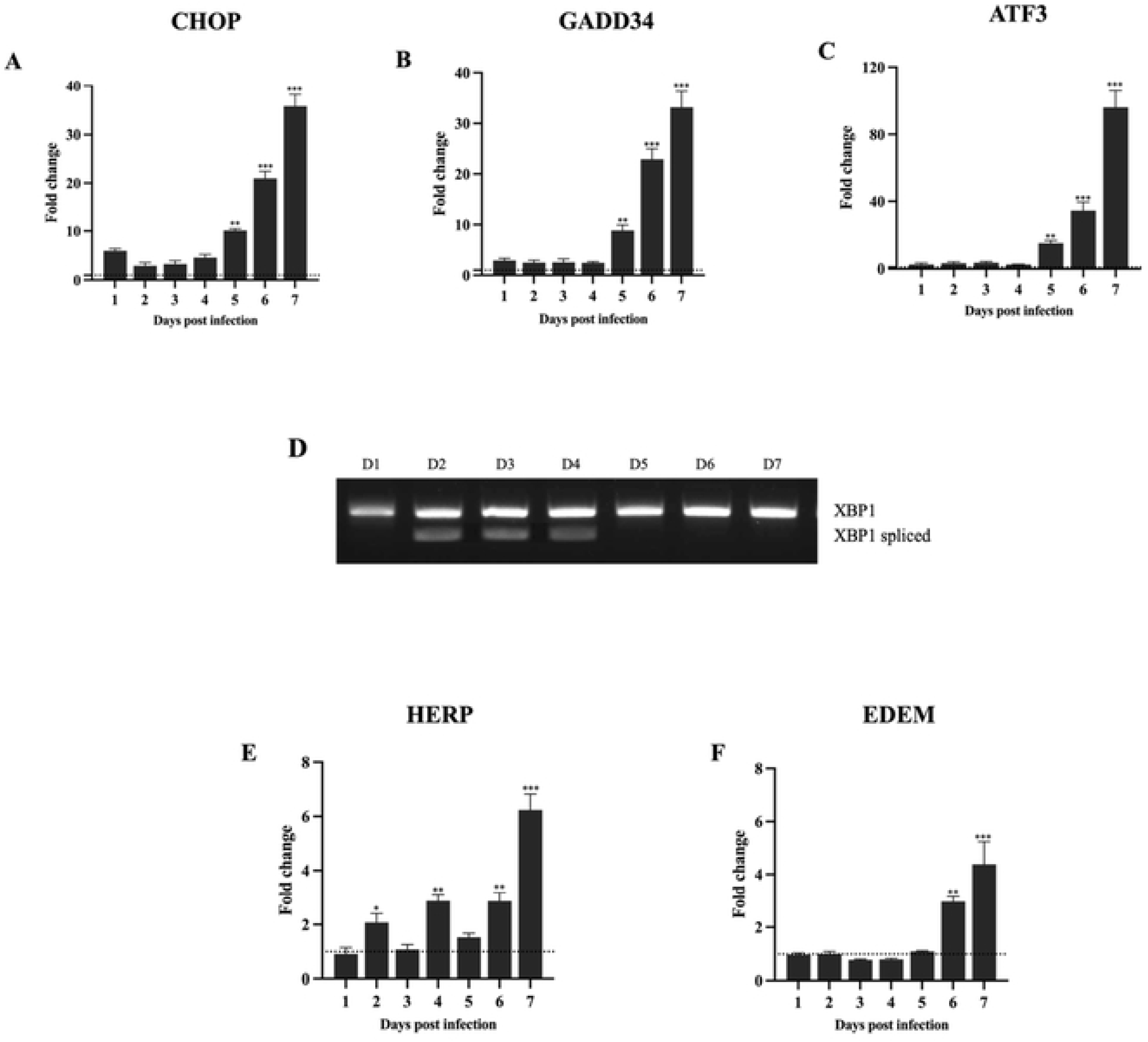
TMUV infection activates the unfolded protein response (UPR) in human lung epithelial A549 cells. A549 cells were infected with TMUVic-C2 at a MOI of 0.1, and UPR-associated gene expression was monitored daily from 1 to 7 dpi by either quantitative RT-qPCR or conventional PCR. (A) Fold change of CHOP. (B, C) Evaluation of modulation of the downstream feedback regulator genes of the PERK-eIF2α pathway: (B) GADD34 and (C) ATF3. (D) Visualization of the splicing of XBP1 mRNA regulator in a 2.5% agarose gel electrophoresis. Lanes D1–D7: Samples collected at 1 to 7 dpi., respectively (E, F) modulation of the ERAD associated gene HERP (E) and EDEM (F). Data represent mean ± SEM of three independent experiments. Statistical significance was determined using one-way ANOVA with post hoc tests *\*p < 0.05, **p < 0.01, ***p < 0.001* compared to fold change of 1 dpi.

To assess the activation of the UPR-IRE1 arm by TMUV infection, we monitored the splicing of XBP1 mRNA into XBP1s mRNA, which can trigger the downstream activation of ERAD-associated genes. An increase in XBP1s mRNA was observed from 2 to 4 dpi (Fig 7D) revealing the effective activation of the IRE1-mediated signaling pathway. Finally, as showed in Fig 7E and 7F, HERP and EDEM gene, two downstream effectors involved in ER-associated degradation (ERAD), were significantly upregulated along the time course. These data suggest that multiple mechanisms of the UPR are activated following TMUV infection in human lung epithelial cells. These findings demonstrate that TMUV infection can trigger ER stress and activates all three canonical UPR pathways (PERK, IRE1, and ATF6) in A549 cells, potentially contributing to host antiviral responses.

## Discussion

Initially reported in Malaysia, the Tembusu virus, an *Orthoflavivirus* highly pathogenic for birds, has been reported in several countries in Asia including China, Thailand, Taiwan, and Vietnam (4, 7, 8, 35). This virus is now considered a zoonotic risk since viral RNA and antibodies have been identified in poultry farm workers and human populations. (20, 21, 36, 37). Although TMUV initially identified as a mosquito-borne pathogen, accumulating evidence suggests that TMUV could adapt to vertebrate hosts through alternative transmission routes, including airborne transmission via the respiratory tract (13, 38). Nevertheless, the molecular basis of virus-host interactions with mammals, particularly with humans, has not yet been fully characterized. However, it is essential to study cellular responses to TMUV infection, as this can provide a better understanding of how the virus modulates the response to infection, thereby improving our understanding of its pathogenesis and enabling the development of strategies to limit viral transmission and zoonotic risk. Here, we report that human lung epithelial cells are highly permissive to TMUV infection, presenting a strong antiviral activity and showing efficient viral replication compared to *Aedes albopictus* mosquito cells. However, this permissiveness depends on the genetic background of the TMUV. These findings may support the hypothesis that TMUV, by infecting human lung epithelial cells, could have the potential for zoonotic transmission via the airborne route, which should be taken into consideration for public health.

As previously described, Clusters 1 and 2 of TMUV represent the major circulating lineages, raising concerns regarding their capacity to infect and adapt to mammalian cells (12, 19, 20). However, Cluster 3 has been reported with increasing frequency in recent years (15). In order to investigate the replication capacity of representative strains from Cluster 1 and Cluster 2 in human lung epithelial A549 cells. We decided to produce two clonal strains of TMUV using reverse genetic engineering. This term encompasses various approaches described in detail by Aubry *et al.* in 2015 (31) and widely used with *Orthoflaviviruses* such as dengue virus and West Nile virus. The production of infectious clones by CPER has already been used successfully to generate JEV and chimeric clones of ZIKV but also with SARS-Cov-2 (29, 39). Previous studies on TMUV used viruses isolated from infected vertebrate hosts or mosquito vectors carrying the virus and collected from the field. However, this involves using a genetically diverse pool of viruses, as genetic drift phenomena of varying degrees occur during the replication course of an *Orthoflavivirus* in the vector or in the vertebrate host (40). Here, we described for the first time the use of the CPER technique to generate infectious clones of TMUV. This simple yet incredibly effective technique enabled us to generate two infectious clones with a well-known sequence. This technique could subsequently be used to generate chimeric or mutant TMUV strains with a known genomic sequence in order to study viral or pathogenic specificities linked to genotypes.

Our results revealed striking differences between the two clusters with the TMUV cluster 2 strain exhibiting a robust replication in human lung cells. In a previous study, Ruangrung *et al.* demonstrated the sensitivity of different human cell lines to TMUV. Human hepatocyte and neuronal cells were found to be permissive to the virus, whereas human lung epithelial cells were almost completely non-permissive (21). Unfortunately, the information provided in the publication is too limited to determine the cluster of the strain used. However, based on the reported collection time and the geographical origin of the viral sample used by Ruangrung *et al.*, we hypothesized that this virus may be classified within the cluster 2. Overall, the investigations support a clear difference in host cell tropism, suggesting that some strains belonging to the cluster 1 could not replicate in human respiratory epithelial cells in contrast to certain strains of the cluster 2, including the one used in our study. In light of previous work in ducks (13) and in human (21), this capacity of the cluster 2 strains may reflect genetic adaptations that facilitate airborne transmission in avian and mammalian hosts. Previous studies have shown that variations in viral structural and non-structural proteins can influence cell tropism and replication efficiency in vertebrate hosts (41). Specifically for TMUV, the E protein and the 3′UTR were found to contribute to differences in infectivity and transmission between the TMUV MM1775 and TMUV CQW1 strains in mosquitoes, with stem-loop I of the TMUV 3′UTR responsible for viral host specificity and the pathogenicity of the virus in mice (42, 43). Our findings are consistent with these reports, supporting that, while initially described as a mosquito-borne virus, specific genetic lineages of TMUV may be better adapted to replicate in mammalian cells, potentially increasing their zoonotic potential (20, 21, 44). However, more investigation is needed to assess the exact involvement of viral proteins or regions of the viral genome in the pathogenicity of the TMUV, but also monitoring its evolutionary pathway and host adaptation mechanisms.

Although previous studies have demonstrated an effective replication of TMUV cluster 1, 2 and 3 in *Ae. albopictus-*derived cells (C6/36) (45, 46), variations in viral strain infectivity have been observed, as the TMUV cluster 2 CQW1 strain that exhibited significantly lower infectivity in replication kinetic in *Ae. albopictus-*derived cells compared to the TMUV MM1775 strain (46). *Culex spp*. have been proposed as the primary natural vectors of TMUV transmission (47), and TMUV has been repeatedly isolated from various *Culex* mosquitoes in the field (48), whereas no natural isolations have been reported from Aedes mosquitoes showing a replication more efficient in the first one than the latter genus. In our study, the TMUV cluster 2 strain also presents a limited replication in *Ae. albopictus*-derived cells that we may expect according to the previous reports.

In our study, the reduced replication of TMUV cluster 2 in *Aedes albopictus*-derived cells may reflect this inherently lower *Aedes* mosquito competence associated with TMUV. However, Sri-in *et al.* showed that the TMUV cluster 2 strain also multiplied less easily than TMUV cluster 1 strain in *Culex tritaeniorhynchus* mosquito (49). We therefore suggest that the low replication of the virus we used in C6/36 cells is not solely due to cells derived from an inappropriate vector, but rather to the adaptation of certain TMUV strains to vectors and hosts. Taken together, our findings suggest an inability of the some TMUV cluster 2 strains to replicate in mosquito while these strains have adapted to vertebrate host. This supports the hypothesis that genetic variation among TMUV clusters can influence both host cell tropism and vector competence, with strains from cluster 2 that may be less suited to efficient transmission by mosquitoes. Nonetheless, further investigation, as well in cells as *in vivo* mosquito, is required to substantiate this hypothesis.

Despite investigations into the immune responses of ducks and chickens to TMUV infection, the innate immune mechanisms to TMUV infection in humans remains poorly understood (26, 50). Our results demonstrate that TMUV infection activates a broad and dynamic innate immune response in human lung epithelial cells. The observed temporal regulation of viral RNA sensors, downstream transcription factors, and effector molecules indicates a tightly host defense mechanism against TMUV. Among RLRs family, RIG-I showed a rapid and transient hyperactivation after infection in A549 cells. This rapid biphasic pattern of activation followed by a reduction is consistent with innate immune responses observed in other viral systems, as Thoresen *et al.* reported in human epithelial cells RIG-I signaling is initiated within one hour of RNA stimulation, peaks between ∼3-6 h, and then declines by 12-24 h (51). Similarly, in influenza B virus infection, early induction of RIG-I-dependent IFN and ISGs is robust at early time points but then tapers off (52). In contrast, MDA5 exhibited a sustained induction. Silencing MDA5 sensor significantly reduced IFN-α/β, IRF3, and IRF7 expression and enhances TMUV while the overexpression of MDA5 strongly potentiated type I interferon and ISG induction, leading to a marked reduction in viral RNA loads. These results align with the canonical functions of MDA5 as viral RNA sensors, which converge on IRF3/IRF7 and NF-κB signaling to initiate antiviral immunity (53). The complementary pattern of RLRs activation is consistent with previous findings of WNV infection, suggesting that PAMPs from early time points after WNV infection are preferentially sensed by RIG-I, leading to an early deficit in IFN-β mRNA induction in RIG-I deficient cells, whereas MDA5 plays a specific role in the later phase of amplification of IFN-β signaling (54). A similar distribution of the response between RIG-I and MDA5 has been observed in DENV and WNV, where both receptors are essential for a robust induction of type I interferon responses (54, 55). The role of MDA5 in controlling flavivirus infection has been previously reported. Thus, in mouse models infected with WNV, MDA5 deficiency resulted in impaired interferon induction and enhanced viral replication, underscoring its role as a critical viral RNA sensor and MDA5 activation is required for efficient interferon responses and suppression of viral replication (53). Our findings extend these observations to TMUV, showing that MDA5-dependent sensing represents a conserved antiviral mechanism across mosquito-borne flaviviruses. Although the immune system is different in birds, recognition of TMUV’s PAMPs and activation of RLRs, particularly LGP2, RIG-I, and MDA5, have previously been also documented in a similar manner in ducks (56).

The endosomal RNA sensors TLRs family, TLR3 and TLR7 were also robustly induced, with TLR3 showing the most striking upregulation. TLR3 is well recognized as a potent sensor of viral double-stranded RNA replication intermediates, inhibition of its expression in chicken DEF cells has been shown to enhance TMUV replication and its activation has been implicated in restricting WNV replication in epithelial and neuronal cells (57, 58). The weaker induction of TLR7 may reflect recognition of single-stranded RNA species generated during TMUV infection, consistent with its role in flavivirus sensing (59). Overexpression of TLR3 appears to significantly enhance the upregulation of type I interferon and ISG induction, which may contribute to a reduction in viral replication. These findings align with the known activity of TLRs, which converge on IRF3/IRF7 and NF-κB signaling pathways to initiate antiviral immunity, though further validation is necessary (58). Importantly, our findings show that loss of TLR3 expression during TMUV infection markedly impaired IFN responses and increased viral burden, underscoring its critical role in epithelial antiviral defense. While RIG-I and TLR7 were also activated during infection, their contributions appeared less prominent compared to MDA5 and TLR3. Overexpression of RIG-I modestly enhanced interferon induction, aligning with its early role in viral sensing whereas TLR7 exhibited only limited effects (54). This may be attributed to its low expression in epithelial cells including A549 cells, in contrast to its higher expression in plasmacytoid dendritic cells (pDCs) and certain macrophage subsets (59). Taken together, our findings highlight a model in which TMUV activates a complex network of RLR- and TLR-dependent sensing pathways, with MDA5 and TLR3 functioning as central nodes. Their activation leads to strong type I IFN and ISG responses, restricting viral replication at epithelial barriers. These findings enhance our understanding of TMUV pathogenesis and highlight the critical role of innate immune signaling in influencing viral evolution and zoonotic potential.

PRR activation converged on the transcription factors IRF3, IRF7, and NF-κB, which were significantly upregulated, correlating with strong induction of type I interferons (60). IFN-β exhibited an early and robust increase, while IFN-α induction was more modest and delayed. This temporal pattern resembles the innate immune response described during WNV infection, IFN-β is identified as an early and critical response, while IFN-α is described as a later-expressed, amplifying cytokine where IFN-β dominates is also plausible given the early IFN-β responses described for ZIKV in human lung epithelial cells (53, 61). As expected, interferon induction is associated with an upregulation of ISGs. The pronounced induction of Mx1, PKR, and OAS1 is indicative of activation of the JAK–STAT signaling pathway, a well-established hallmark of host antiviral immunity (62). This observation is consistent with previous transcriptomic study using duck embryonic fibroblasts infected with DTMUV (63). In addition to antiviral signaling, TMUV infection triggered strong pro-inflammatory responses, with TNF-α, CXCL8, and CXCL10 showing substantial induction. CXCL10 is one of the cytokines, along with IL-6, IL-8, and IL-17, that show the highest induction amplitudes during infections with highly pathogenic viruses such as DENV, ZIKV, and WNV (64). While these inflammatory mediators contribute to viral clearance, their excessive induction has been linked to immunopathology in flavivirus infections (56). Altogether, our observations may reflect that A549 cells had reached an established antiviral state, characterized by high ISG expression and suppression of further interferon signaling to avoid excessive inflammation. Similar transient dynamics have been described in flavivirus infections, following the early induction of phosphorylated STAT1 and type I IFNs, their expression and activity rapidly decrease due to negative feedback regulation through ISGs SOCS1/3 and USP18, which blocks STAT phosphorylation and dampen IFN signaling (65, 66). Alternatively, TMUV may actively antagonize host immunity at later stages, as observed in DENV and ZIKV infections, where viral NS proteins as well as subgenomic flaviviral RNAs (sfRNAs) inhibit JAK-STAT signaling and antagonize innate immunity in order to evade a sustained antiviral cellular response (67, 68). This inhibition effect of the viral NS protein in human needs to be more investigated for TMUV in future.

The UPR, a critical cellular defense mechanism against viral infections, is mediated by three canonical signaling pathways: PERK, IRE1 and ATF6. These arms aim to restore ER homeostasis by reducing protein load, enhancing protein folding capacity, and promoting the degradation of misfolded proteins. However, prolonged or severe ER stress can lead to apoptosis and autophagy (34). Consistent with activation of the PERK pathway, we observed a strong induction of the pro-apoptotic transcription factor CHOP expression. The upregulation of CHOP is a hallmark of ER stress-mediated apoptosis in *Orthoflavivrus* including TMUV (69–71), suggesting that TMUV infection drives programmed cell death in A549 cells (28). In parallel, the IRE1 pathway was activated during TMUV infection, as evidenced by increased expression of HERP and EDEM both of which are ERAD downstream effectors. The late and robust induction of these UPR markers indicates that TMUV-associated ER stress is not immediate but accumulates during viral replication. Furthermore, splicing of XBP1 mRNA was clearly and rapidly detected suggesting that IRE1 activation may function as an early sentinel response to infection, preceding the more delayed upregulation of downstream stress-related genes. An identical splicing pattern was reported by Zhao *et al*., with a slightly shorter time frame; however, these observations were made on BHK21 cells infected with a TMUV cluster 2 strain (28). Similar temporal differences in UPR activation have been reported for other orthoflaviviruses, where early XBP1 splicing primes ER-associated degradation, while later activation of CHOP might reflects the onset of apoptosis and feedback regulation, confirming the engagement of IRE1-mediated signaling, which plays a dual role in restoring ER function and regulating innate antiviral responses (33, 69).

We also observed marked the upregulation of GADD34 and ATF3 at late stage of infection, indicative of an activation of the PERK-eIF2α feedback loop and stress-associated transcriptional reprogramming, respectively. In previous report, GADD34 was shown to be activated by TMUV infection in BHK21 since 24 hpi (28). ATF3 is known to integrate signals from multiple stress pathways and has been implicated in apoptosis, cytokine regulation, and immune modulation during viral infections (72). Further highlighting that TMUV-induced ER stress culminates at later stages of infection. The strong late-phase induction of ATF3 suggests that TMUV infection imposes sustained ER stress, driving transcriptional reprogramming toward apoptosis and immune modulation. Together, our data indicate that TMUV infection in A549 cells triggers a biphasic UPR response; an early IRE1-XBP1 activation during initial replication, followed by late activation of CHOP, HERP, EDEM, GADD34, and ATF3 coinciding with peak viral replication. This sequential activation suggests that TMUV initially engages the UPR to accommodate increased protein folding and replication demands, but prolonged infection shifts the response toward apoptosis and stress-induced cell death. Such dynamics have been similarly observed in JEV and WNV, where late UPR activation correlates with cytopathic effects and viral pathogenesis (33). However, further investigation will be necessary to evaluate the effective implementation of apoptosis and autophagy in response to infection of human cells by TMUV cluster 2.

Overall, our findings emphasize that TMUV infection in human lung epithelial cells elicits both robust viral replication and innate immune activation, while simultaneously inducing significant ER stress and UPR signaling pathway activation. These observations suggest that specific strains of TMUV may have adapted to mammalian hosts, raising important questions about the pathogenesis of the virus in humans. It will be essential to continue research into the molecular mechanisms involved in infection, particularly the impact of UPR pathway activation and the role of innate immunity in the development of infection in humans. Further research into the TMUV pathogenesis in humans is crucial to assess potential risks to human health and inform strategies for prevention and control. The adaptation to humans and airborne transmission mode encourages us to continue research into the epidemiological potential of TMUV in human populations.

## Acknowledgements

The authors would like to express their sincere gratitude Associate Professor Dr. Hatairat Leardsamran for her valuable technical suggestions regarding the experiments, as well as for her financial support and mentorship to Kachaporn Jintana. We would like to thank Dr. Sebastien Nisole for kindly providing us the A549 cells used in this study.

## Supporting information

**S1 Fig. Validation of MDA5, RIG-I, TLR3 and TLR7 overexpression in A549 cells.** A549 cells were transfected with plasmid encode targeting MDA5, RIG-I, TLR3 or TLR7. At 24, 48, and 72 hpi, overexpression efficiency was confirmed by qRT-PCR. Data represent mean ± SD of three independent experiments. Statistical significance was determined using one-way ANOVA with post hoc tests (*\*p < 0.05, **p < 0.01, ***p < 0.001* compared to uninfected cell control).

**S2 Fig. Validation of MDA5 and TLR3 knockdown in A549 cells.** A549 cells were transfected with siRNA targeting MDA5 or TLR3. At 24, 48, and 72 hpi., knockdown efficiency was confirmed by qRT-PCR. Statistical significance was determined using one-way ANOVA with post hoc tests (*\*p < 0.05, **p < 0.01, ***p < 0.001* compared to cells transfected with control siRNA).

**S1 Table. Primer sequences for viral infectious clones’ generation and linker generation.**

**S2 Table. Primer and probe sequences for TMUV viral genome quantification.**

**S3 Table. List of primer used for RT-qPCR.**

## Notes

### Competing Interest Statement

The authors have declared no competing interest.

